# LATS2 modulates synaptic organization and function by regulating actin dynamics

**DOI:** 10.64898/2026.10.16.758881

**Authors:** Michael Wigerius, Parabhjot K Labana, Antonios Diab, Stefan Krueger, James P. Fawcett

**Affiliations:** Department of Pharmacology, Dalhousie University, Halifax NS, B3H 4R2; Department of Physiology and Biophysics, Dalhousie University, Halifax NS, B3H 4R2; Department of Surgery, Dalhousie University, Halifax NS, B3H 4R2

## Abstract

Recent work has highlighted the importance of core Hippo pathway kinases in cycling cells. Many of these kinases are also expressed in post-mitotic neurons, where they have been linked to synaptic function, and more recently a number have been shown to be dysregulated in neurodegenerative diseases such as Alzheimer’s disease. Despite this, little is known about the specific role or subcellular localization of the Large Tumor Suppressor kinases, LATS1 and LATS2, in postmitotic neurons. Previously, we demonstrated that LATS1 plays an important role in synapse development; however, the role of LATS2 remains poorly understood. Here, we examined the distribution and function of LATS2 and found that it is highly expressed in CA3/CA4 neurons of the hippocampus and localized throughout dendrites and dendritic spines. In contrast to LATS1, LATS2 expression remains stable throughout neuronal development. Furthermore, ectopic expression of LATS2 induces kinase-dependent structural spine defects, alters actin dynamics and reduces activity-dependent synaptic SEP-GluR1 fluorescence. Collectively, these findings reveal a unique distribution LATS2 in the hippocampus and demonstrate that LATS2 directly regulates actin dynamics and GluR1-containing AMPAR trafficking.

**Significance Statement:** The Hippo kinases LATS1 and LATS2 have been studied in proliferating cells, yet their function in post-mitotic neurons remain poorly understood. Here, we identify a distinct role for the LATS2 kinase in hippocampal neurons, where we find it is enriched in CA3/CA4 regions and localized to dendrites and dendritic spines. Unlike LATS1, we find LATS2 is maintained throughout neuronal maturation. We find LATS2 regulates dendritic spine structure, actin dynamics, and the trafficking of GLuR1-containing AMPA receptors in a kinase-dependent manner. These findings establish LATS2 as a novel regulator of synaptic structure and function and provide further evidence that Hippo pathway kinases play a role in synaptic morphology and function.

## Introduction

The NDR/LATS proteins, large tumor suppressor 1 and 2 (LATS1 and LATS2), are serine-threonine kinases that function as core components of the hippo pathway, which is activated in response to various stimuli, including cell adhesion and mechanical tension (Meng et al., 2016). They are closely related to the Ndr1/2 kinases which have been reported to be involved in regulating different aspects of neuronal development (Santos et al., 2023). Both LATS1 and LATS2 are ubiquitously expressed in cells where they exert tumour suppressor activity but are present at lower levels in cancers due to their downregulation or catalytic inactivation (Dey et al., 2020). Their main targets are the transcription co-activators YAP and TAZ, that function to drive proliferation by potentiating transcriptional activity in the nucleus. Both LATS1 and LATS2 inhibit YAP and TAZ through phosphorylation of key serine residues causing their cytoplasmic retention, thereby regulating tissue growth.

In some contexts, LATS1 and 2 share redundant functions but studies have revealed functional differences in part due to the networks that control their mRNA expression and subcellular localization (Furth and Aylon, 2017). While both kinases have been primarily studied in immortalized cell systems their activity in postmitotic cells and particularly the nervous system, are only now beginning to emerge (Wigerius et al., 2018; Wigerius et al., 2020). Hippo signaling components including MST1-3 and NDR1/2 (Ultanir et al., 2012; Lee et al., 2013; Tang et al., 2014), along with YAP and its effector TEAD1 (Rojek et al., 2019; Zaltsman et al., 2019; Atterton et al., 2025), have all been implicated in nervous system processes, including development and neurodegenerative disease. We have found that LATS1-dependent phosphorylation regulates association between the scaffolding protein Angiomotin (AMOT-130) and actin filaments during development in primary hippocampal neurons (Wigerius et al., 2018). Yet, very recent reports suggest that LATS2, of the two proteins, is the principal kinase in the brain, with functions independent of LATS1.

Reprogramming of hippocampal gene expression, whether induced by exposure to a novel environment or by epigenetic signaling triggered by the metabolite acetate, alter LATS2 brain expression with downstream effects on learning and memory (Traunmuller et al., 2025; Periandri et al., 2026). Moreover, studies link LATS1 and LATS2 to a-amino-3-hydroxyl-5-methyl-4-isoxazole-propionate receptor (AMPAR) trafficking through its regulation of Kibra (WWC1), a scaffolding protein that is required for synaptic transmission and long-term memory (Makuch et al., 2011; Stepan et al., 2024). However, it remains unclear whether LATS kinases play a direct or indirect role in facilitating these interactions or how it is distributed in neurons.

In mammals, excitatory synapses form on dendritic spines, which are actin-rich protrusions on neighbouring neurons that are critical for structural synaptic plasticity and long-term memory formation (Matus, 2000). Pathological changes in spine structure during formation or development contributes directly to neurological disorders (Penzes and Cahill, 2012). At the tip of the spine lies an electron dense structure, the post-synaptic density (PSD), which contains a specialized network comprising distinct scaffolding proteins, adapters and kinases. The components of the PSD functions to anchor and stabilize AMPARs thereby modulating the balance between synaptic strength and weakening (Huganir and Nicoll, 2013).

In this study we have examined the distribution and functions of LATS2 in primary hippocampal neurons. We found that LATS2 is widely distributed along dendrites and spines and in contrast to LATS1, is steadily expressed in neurons throughout development. Our results furthermore show that overexpression of LATS2 causes structural spine defects that are kinase-dependent, leading to changes in actin dynamics and a reduction in activity-dependent synaptic SEP-GluR1 fluorescence. Collectively, these findings provide insight into the distribution and functions of LATS2 in dissociated neurons as well as an unanticipated effect on GluR1-containing AMPAR trafficking.

## Materials and methods

### Constructs and antibodies

Plasmids with cDNA encoding pCIneoMyc-LATS2, pEGFP-b-Actin, and tdTomato have been described previously (Wigerius et al., 2018; Quinn et al., 2019). The LATS2-ΔC (2160 bp) and LATS2-ΔN (1263 bp) deletion constructs were generated by PCR amplification (Phusion High-Fidelity PCR System; New England Biolabs) and inserted into pCDNA3-2x-Myc using the BamH1 and EcoR1 restriction sites. The point mutation (K697A) in the pCIneoMyc-LATS2-K697A construct was generated by using the Phusion Site-Directed mutagenesis kit (Thermo Scientific). All constructs were verified by sequencing. The following plasmids were obtained from Addgene: pcDNA-HA-LATS2-KR (#33100; a gift from Kun-liang Guan; Westlake University, Zhejiang, China), pCI-Sep-GluR1 (#24000; a gift from Robert Malinow; University of California, San Diego, USA). Primary antibodies were as follows: LATS2 (1:500; NBP3-03913; Novus Biologicals), PSD-95 (1:500; ab2723; Abcam), Bassoon (1:400; 140-011; Synaptic Systems), myc-9E10 (1:500; sc-40; Santa Cruz Biotechnology), HA (1:2000; H3663; Sigma-Aldrich), GFP (1:5000; ab290; Abcam), pS175-AMOT (1:400; ABS1045; EMD Millipore), b-Actin (1:2000; MA1-40; Invitrogen). Horseradish peroxidase (HRP)-conjugated secondary antibodies were used follows: IgG-HRP mouse (1:4000; sc-2768, Santa Cruz Biotechnology) and IgG-HRP rabbit (1:4000; 31466; Thermo Fisher Scientific). For immunostaining Alexa Fluor 488 anti–mouse, Alexa Fluor 488 anti– rabbit, Alexa Fluor 594 anti–mouse, Alexa Fluor 594 anti–rabbit (1:1000; A-11029, A-11034, A-11032, and A-11037, respectively (Thermo Fisher Scientific), were used.

### Cell culture and transfection

Dissociated cultures of primary hippocampal neurons were prepared from E18 Sprauge-Dawley rat embryos as described previously (Richier et al., 2010). Experiments were performed in 11-17 DIV hippocampal neurons. For biochemical analysis neurons grown for 22 days in vitro (DIV) were solubilized in mild lysis buffer containing: (20 mM Tris [pH 8.0], 137 mM NaCl, 10 % glycerol, 1 % NP-40), supplemented with protease inhibitors and centrifuged at 13 800 g for 20 minutes at 4°C. Supernatant fractions were analyzed by Western blotting. HEK293T and COS-7 cells were maintained at 37°C and 5 % CO_2_ in DMEM supplemented with 10 % FBS, 10 U/ml penicillin and 10 µg streptomycin. For neuron transfections, the calcium phosphate method was used as previously described (Richier et al., 2010). HEK293T and COS-7 cells were transfected with polyethylenimine (PEI) by mixing 3 µg of each DNA construct with PEI (1 mg/ml) at a ratio of 4:1 PEI to DNA.

### In vivo phosphorylation assay

For detection of AMOT-130 phosphorylation in vivo, HEK293 cells were transfected as indicated with empty vector, myc-LATS2, myc-LATS2-KA or LATS2-KR using PEI. Twenty-four hours after transfection, cells were lysed in mild lysis buffer, and lysates were cleared by centrifugation. After determination of protein concentrations using the bicinchoninic assay method (Thermo Fisher Scientific), samples were loaded for SDS-PAGE and transferred to nitrocellulose membrane (Bio-Rad). Membranes were probed with pS175-AMOT antibody overnight followed by incubation with secondary HRP-conjugated antibody and visualized by chemiluminescence (Thermo Fisher Scientific). The Western blots were re-probed with the indicated antibodies.

### Immunoprecipitation

For rat brain immunoprecipitations, crude synaptosomal P2 fractions prepared as previously described (Wigerius et al., 2018), were solubilized at RT in lysis buffer and centrifuged at 13 800 g for 45 minutes at 4°C. Lysates were incubated with 1–2 µg of either isotype-matched IgG, anti–myc, or anti–LATS2 antibody at 4°C overnight followed by incubation with protein-A Sepharose beads (GE Healthcare). For analysis of protein interactions in HEK293 cells, GFP-Actin was co-transfected with the indicated myc-tagged LATS2 constructs and immunoprecipitated with anti GFP-antibody followed by protein-A Sepharose beads incubation. The beads were washed three times in lysis buffer, eluted in 3x Laemmli sample buffer. Bound proteins were subjected to Western blotting and visualized as described. Relative protein intensities were quantified by densitometry using the open-source platform Fiji (National Institute of Health, NIH) and visualized by line scan plots.

### Cell stimulation and immunocytochemistry

15-16 DIV neurons (4 days post-transfection) were incubated for 10 minutes in Extra Cellular Solution (ECS) containing:(150 mM NaCl, 2 mM CaCl2, 5 mM KCl, 10 mM Hepes, 30 mM Glucose, Tetrodotoxin (TTX) 1 µM, and Picrotoxin (PTX) 30 µM). To induce chemical activity in vitro, neurons were incubated without or with 0.3 mM glycine in ECS at room temperature (RT) for 4 minutes, and then for another 10 and 20 minutes in ECS without glycine. Neurons were fixed in 4 % PFA/4 % sucrose/1x PBS solution for 15-20 minutes at RT and permeabilized in 0.1 % Triton-X100/1x PBS solution for 10 minutes. For detection of endogenous LATS2, neurons were fixed for 3 minutes in 4 % PFA/4 % sucrose/1x PBS followed by fixation for another 10 minutes in ice-cold methanol. Fixed cells were washed extensively in 1x PBS and then blocked in 5 % BSA in PBS with 0.1 % Triton-X100 for 2-3 hours at RT. After blocking, cells were incubated with primary antibodies in 5 % BSA/0.1 % Triton-X100/1x PBS solution overnight at RT. Then, cells were washed in 1x PBS three times for 5 minutes each, incubated with secondary antibodies in 5 % BSA/1x PBS solution for 1-2 hours at RT, washed in 1x PBS, and mounted on cover slips with fluoromount (Sigma-Aldrich).

### FRAP

Frap experiments were performed as previously described (Wigerius et al., 2018; Diab et al., 2023). Briefly, imaging was conducted on a spinning-disk microscope (Zeiss) using a 63 x 1.4 oil immersion lens with a stage incubator (37°C and 5 % CO_2_). Dendritic spines on hippocampal neurons expressing either GFP-Actin alone or together with WT or mutant LATS2 for 3-4 days were photobleached and imaged. To monitor Actin turnover, a ROI with a diameter of 1 ∼ µm was photobleached at full laser power (100 % power and 100 % transmission) for 2.5 seconds. Fluorescence recovery was monitored by automatic scanning of the whole cell in 0.2-second intervals at low laser power. During image processing the mean fluorescence intensity of an un-transfected region was measured as background and subtracted from each frame using the Fiji platform. The corrected intensity values were then fitted by nonlinear regression to a one-phase exponential association model in Prism (GraphPad software). The mobile fraction of GFP-Actin was approximated by a photobleach correction equation, as described previously (Feder et al., 1996; Lippincott-Schwartz et al., 1999), and results were presented as percentage by bar graphs made with Prism.

### Quantitative PCR (qPCR)

Total RNA was isolated from embryo and adult rat hippocampi using TRIzol reagent and reverse-transcribed with SuperScript III Reverse Transcriptase according to the manufacturers’ instructions (Invitrogen Life Technologies). LATS2 expression levels were analyzed with qPCR using the SsoFast EvaGreen Supermix Kit as per manufacturers’ instructions (Bio-Rad Laboratories). The threshold cycle (C_T_) value for cyclophilin A was used to normalize LATS2 transcript levels using the DDC_T_-method as previously described (Livak and Schmittgen, 2001).

The following primers were used for qPCR:

*LATS2_1L*: ’5-CGAGCAAGACAGGAATGACA-3’

*LATS2-1R*: ’5-GCTGACCTTCTGCTGGTAGG-3’

*Cyclophilin A-5A*: 5’-GAGCTGTTTGCAGACAAAGTTC-3’

*Cyclophilin A-3A*: 5-CCCTGGCACATGAATCCTGG-3’

### Analysis of Human Protein and Allen Brain Atlas datasets

To gain insight into LATS2 mRNA expression in the brain, we used the Human Brain Altas single-nuclei RNA sequencing (snRNAseq) dataset reported by Siletti et al (Siletti et al., 2023). The dataset comprises ∼ 3 million cells across 11 brain regions obtained from adult postmortem donors. We analyzed 21 clusters from this dataset, presented as a histogram in the current study, using the same naming system of cells and brain regions as defined in <u>The Human Protein Atlas</u>. The in-situ hybridization image and corresponding LATS2 expression map in adult mouse brain was obtained from the Allen Mouse Brain Atlas (https://mouse.brain-map.org/experiment/show/73512206).

### Confocal image acquisition and analysis

Microscopy and post analysis of neurons was performed as previously described (Wigerius et al., 2018). Briefly, images were acquired as z-stack series taken at 0.3-µm step intervals with an LSM710 confocal scanning microscope using 40x 0.7 NA and 63x 1.4 NA oil immersion lenses and collected with a digital Axio-cam camera controlled by ZEN software (ZEISS). Parameters were the same across all experiments, and 10-15 neurons were analyzed for each experimental condition. Quantitative analysis of fluorescent intensity distribution of endogenous proteins was performed in Fiji by outlining puncta on spines and dendritic shafts using regions of interest (ROIs) that was overlaid on 100 µm dendritic segments. Results were presented as the total integrated fluorescence intensity of each protein. For correlation of LATS2, myc-LATS2, HA-LATS2-KR clusters overlapping with synaptic markers, dendritic ROIs were generated using thresholding and the wand tool on a single channel. The ROIs were then applied to the corresponding second channel and areas outside the ROIs were cleared to exclude background signal from un-transfected neurons. The processed channels were merged and the overlap in pixel intensity on dendrites and synapses was described by the Pearson’s correlation coefficient calculated in Prism. Differences between Pearson coefficients were determined using Fisher’s r-to-z transformation. Puncta overlap was analyzed by the Fiji ‘Colocalization’ plugin yielding a pixel map of colocalized clusters. To analyze spine enrichment of myc-LATS2, myc-LATS2-ΔC, and myc-LATS2-ΔN, spines were outlined as described above, and the spine enrichment coefficient (E_f_) was calculated by measuring the ratio of spine-to-shaft fluorescence intensity of the myc-tag signal to tdT. All images were collected from at least two independent experiments per conditions and analyzed with the experimenter blinded to the conditions.

### Data handling and statistical analysis

Statistical analysis was performed in Prism (10) software, and all data sets were checked for Gaussian normality distribution. Data testing of two groups was performed using an unpaired Student’s t-test with Welch’s correction. Multiple group analysis was performed using one-way or two-way Anova followed by Tukey’s or Sidak’s test, respectively, to account for multiple comparisons. Statistical significance was set at P < 0.05. In figures significance levels were *P < 0.05, **P < 0.01, *** P < 0.001 and **** P < 0.0001 as defined in Prism. Quantitative analysis of spine length, area, and density was performed manually in Fiji as described (Wigerius et al., 2018), and each experiment was reproduced at least two times with independent neuronal cultures. Spine morphometric values measured on individual neurons were first averaged per neuron, and means from several neurons were then averaged to obtain the population mean. Measurements in tables are given as means ± SEM. All figures were created in Illustrator using the Creative Cloud platform (Adobe version 30.x).

## Results

LATS1 has been previously implicated in the nervous system, and publicly available data sets indicate that also LATS2 is expressed in neural tissue. We therefore characterized the gene expression profile of *LATS2* by qPCR from samples collected from embryo and adult mouse hippocampi. We found that *LATS2* expression was significantly elevated in adult compared to embryo samples (**Fig. 1A**; \*\**P* < 0.01, unpaired *t*-test with Welch’s correction). To validate the global *LATS2* gene expression profile, we reprocessed publicly available datasets that applied single nucleus RNA sequencing (snRNA-seq) to > 3 million cells sampled across 11 human brain regions (<u>Human Protein Atlas</u>). Based on these datasets, we found that *LATS2* is heterogeneously expressed across brain regions with peak values in CA4 that were several-fold higher compared to other regions (**Fig. 1B**). Additionally, cells in the CA1-3 subregions, specific astrocyte subtypes, and cells in the midbrain and upper rhombic lip, regions primarily involved in early development and cerebellar formation, exhibited intermediate to moderate levels of expression. These observations were further validated using open access in situ hybridization data from the Allen Brain Atlas (<u>Allen Brain Atlas</u>), which showed elevated *LATS2* expression particularly within the CA3 and CA4 regions (**Fig. 1C**). Thus, region-specific *LATS2* expression observed across independent datasets suggest a specialized neuronal role, consistent with LATS2 expression increasing in post-mitotic neurons and reports of increased LATS2 levels in hippocampal subregions following novel environment exposure (Traunmuller et al., 2025).

**Figure 1.**
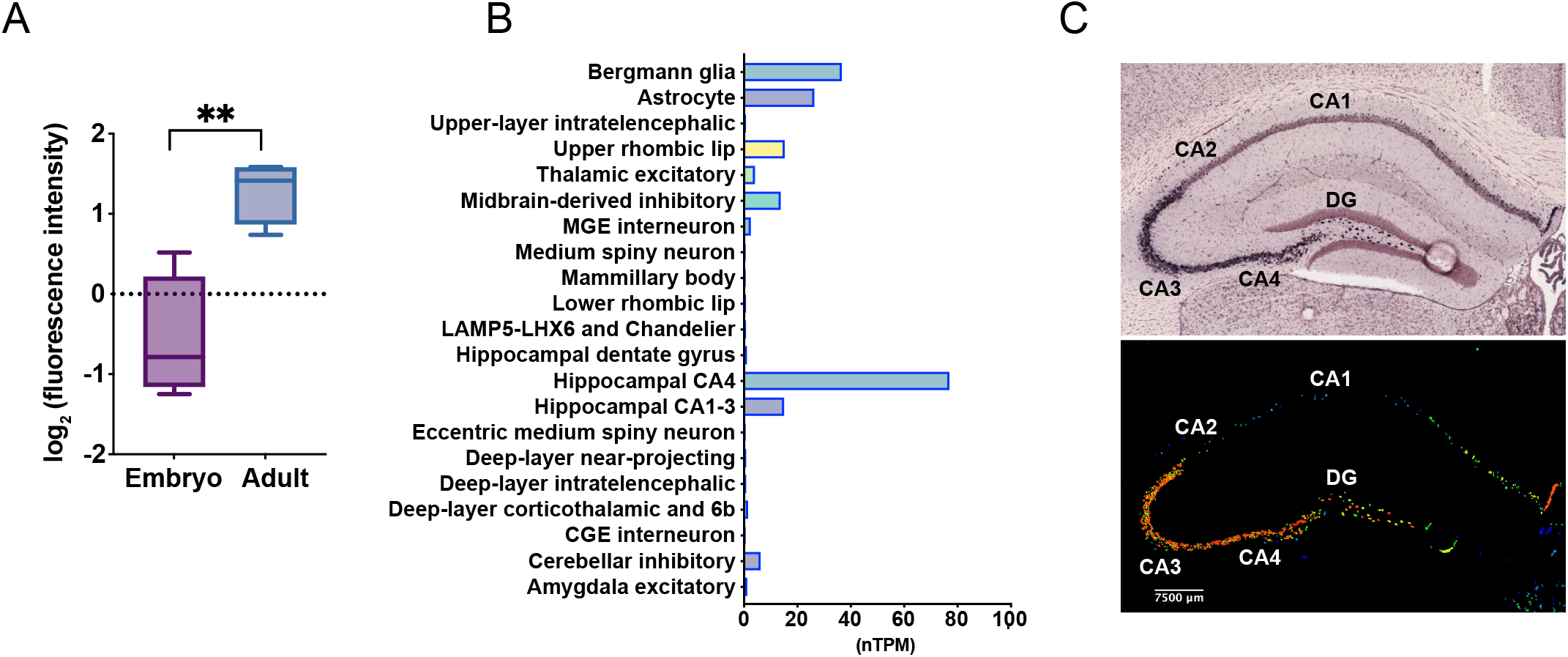
LATS2 gene expression increases during mouse brain development. **A.** Log_2_-transformed plot of the relative LATS2 gene expression in rodent Embryo (E18) versus Adult (P56) hippocampal tissue assessed by RT-qPCR. Expression data are presented as fold change. \*\**P* < 0.01, unpaired *t*-test with Welch’s correction. **B.** Histogram of LATS2 gene expression across different human brain tissues assessed by snRNA-seq shown as normalized Transcripts Per Million (nTPM). Source: Human Protein Atlas. Note elevated expression of LATS2 specifically in CA3/CA4 region neurons. **C**. Zoomed in view of LATS2 expression in the adult (P56) mouse hippocampus. ISH image (top) and corresponding gene expression map (bottom), with CA1-CA4 and DG labelled. Scale bar, 7000 µm. The ISH and expression map images were obtained from the Allen Brain Atlas (https://mouse.brain-map.org/experiment/show/73512206).

Having examined the global gene expression profile, we next characterized the expression of LATS2 in the brain and in cultured hippocampal neurons. Immunoprecipitation of lysate from crude synaptosome fractions showed enrichment of LATS2 in the adult rat brain (**Fig. 2A**). To assess LATS2 in neurons we probed equal amounts of hippocampal whole cell lysate prepared at different developmental stages with anti-LATS2 antibody. Following an increase in expression early in development, LATS2 levels remained uniform across the remaining time points analyzed (**Fig. 2B**). We next stained neurons for endogenous LATS2, which demonstrated a punctate pattern along dendritic shafts and spine heads that overlapped with Bassoon, a key component of the pre-synapse (**Fig. 2C**). Cumulative frequency histograms revealed no difference in mean fluorescence intensity across channels (**Fig. 2D**), suggesting that puncta overlap was independent of overall signal abundance. To quantify the compartment-specific overlap in neurons, we performed Pearson correlation and regression analyses of LATS2 and Bassoon fluorescence intensities along dendrites. We found that LATS2 and Bassoon exhibited a stronger correlation overlap in dendritic shafts compared to spine heads (**Fig. 2F,G**; spines: *r* = 0.718; shafts: *r* = 0.845; \*\**P* = 0.037; *Z* = 2.09, two-tailed Fisher’s r-to-z test). These analyses were performed in mature (DIV14-16) cultures with puncta samples quantified across primary and secondary dendrites, suggesting that the variability in LATS2-Basson puncta overlap may reflect compartment-dependent functional differences.

**Figure 2.**
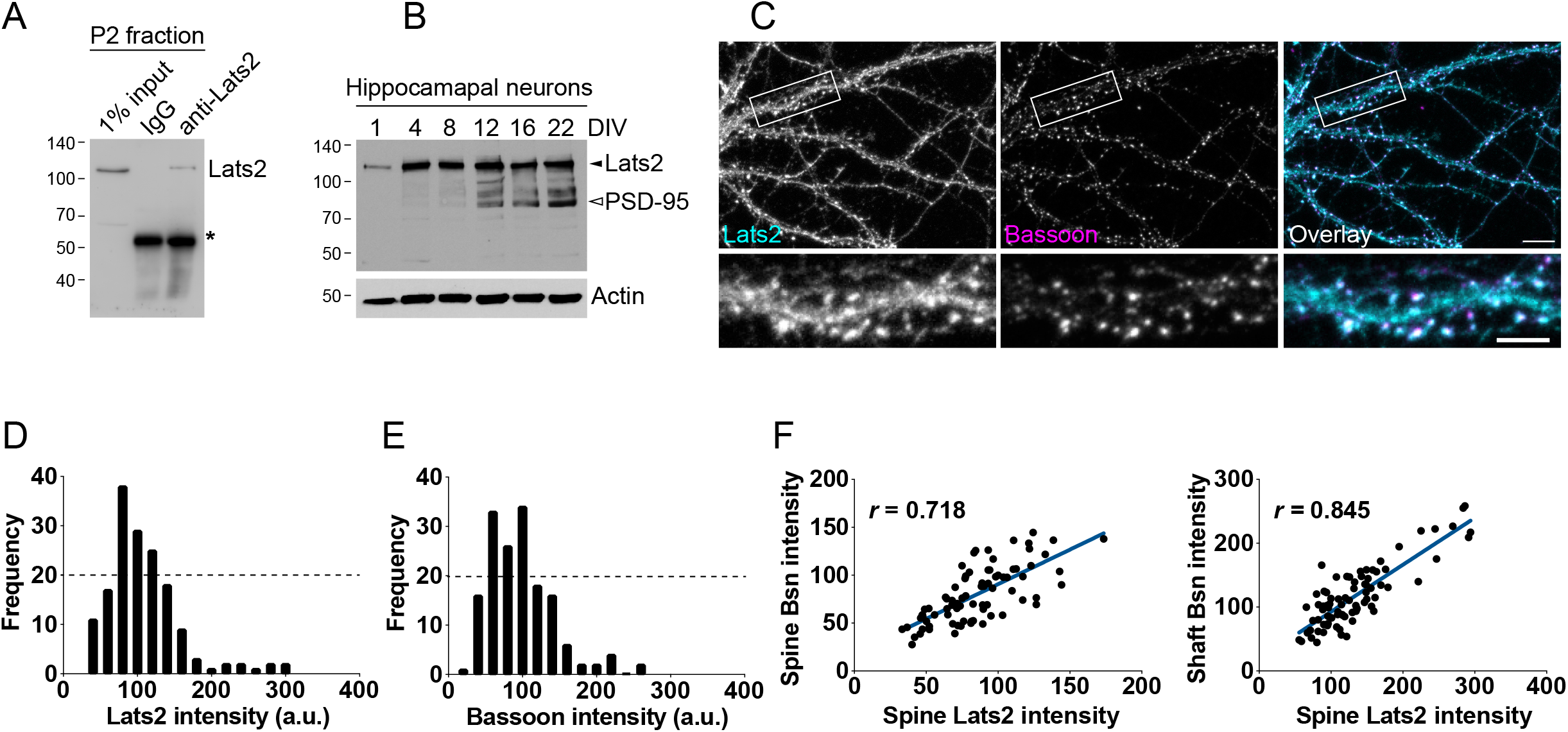
LATS2 expression in dissociated hippocampal neurons. **A.** Immunoprecipitation (IP) of endogenous Last2 from the crude synaptosome fraction (P2) of adult rat brain. IP samples were analyzed by immunoblotting with LATS2 antibody. The asterisk indicates the IgG heavy chain. **B.** Western blot analysis of whole neuron lysate collected at 1, 4, 8, 12, 16, 22 Day In Vitro (DIV), was analyzed with LATS2 antibody. The membrane was sequentially probed for PSD-95 and actin. **C.** Double immunostaining of endogenous LATS2 (cyan) and Bassoon (Bsn, magenta) in hippocampal neurons (DIV17). Scale bars, 10 µm (top); 5 µm (bottom). **D**, **E**. Histogram of LATS2 and Bassoon fluorescence intensity distribution in neurons. Arbitrary units (a.u.). **F**. Pearson’s correlation analysis of LATS2-Bassoon puncta in spines (*r* = 0.718) and in dendritic shafts (*r* = 0.845); \**P* = 0.037; *Z* = 2.09, two-tailed Fisher’s r-to-z test.

To examine whether the kinase activity of LATS2 influence spine structure, we first assessed the ability of LATS2 wild-type (WT) and two kinase-inactive mutants (K697A and K655R) to phosphorylate AMOT-130 at serine 175 in HEK293 cells. Equal amounts of lysate from cells transfected with either myc-LATS2, myc-LATS2-KA or HA-LATS2-KR were analyzed by immunoblotting using a phospho-site specific antibody (pS175). Whereas wild-type LATS2 induced AMOT-130 phosphorylation, both mutants exhibited reduced phosphorylation, consistent with a loss of kinase activity (**Fig. 3A**, **3B**). We next co-transfected the constructs in 11-DIV hippocampal neurons together with GFP-Actin to label dendritic spines and imaged cells 4 days later (**Fig. 3C**). Cumulative frequency analysis showed shifts in distribution of spine length and spine area whereas spine density was unaltered between the groups (**Fig. 3D-F**, **3F**, **3I**). Quantitative analysis revealed that co-expression of GFP-Actin and HA-LATS2-KR induced elongated, filopodia-like, protrusions that were approximately 38 and 48% longer than those observed in cells expressing either only GFP-actin or myc-LATS2 and GFP-actin (**Fig. 3G**; \*\*\*\**P* < 0.0001, one-way Anova with Tukey’s multiple comparison). In contrast, myc-LATS2 expression led to a 30% and 60% increase in spine head area compared to GFP-Actin alone or HA-LATS2-KR, respectively (**Fig. 3H**; \*\**P* < 0.01; \*\*\*\**P* < 0.0001, Anova with Tukey’s multiple comparison). Thus, spine defects characterized by enlarged spine heads and a high number of immature protrusions indicate that the role of LATS2 in spine morphogenesis is kinase activity dependent.

**Figure 3.**
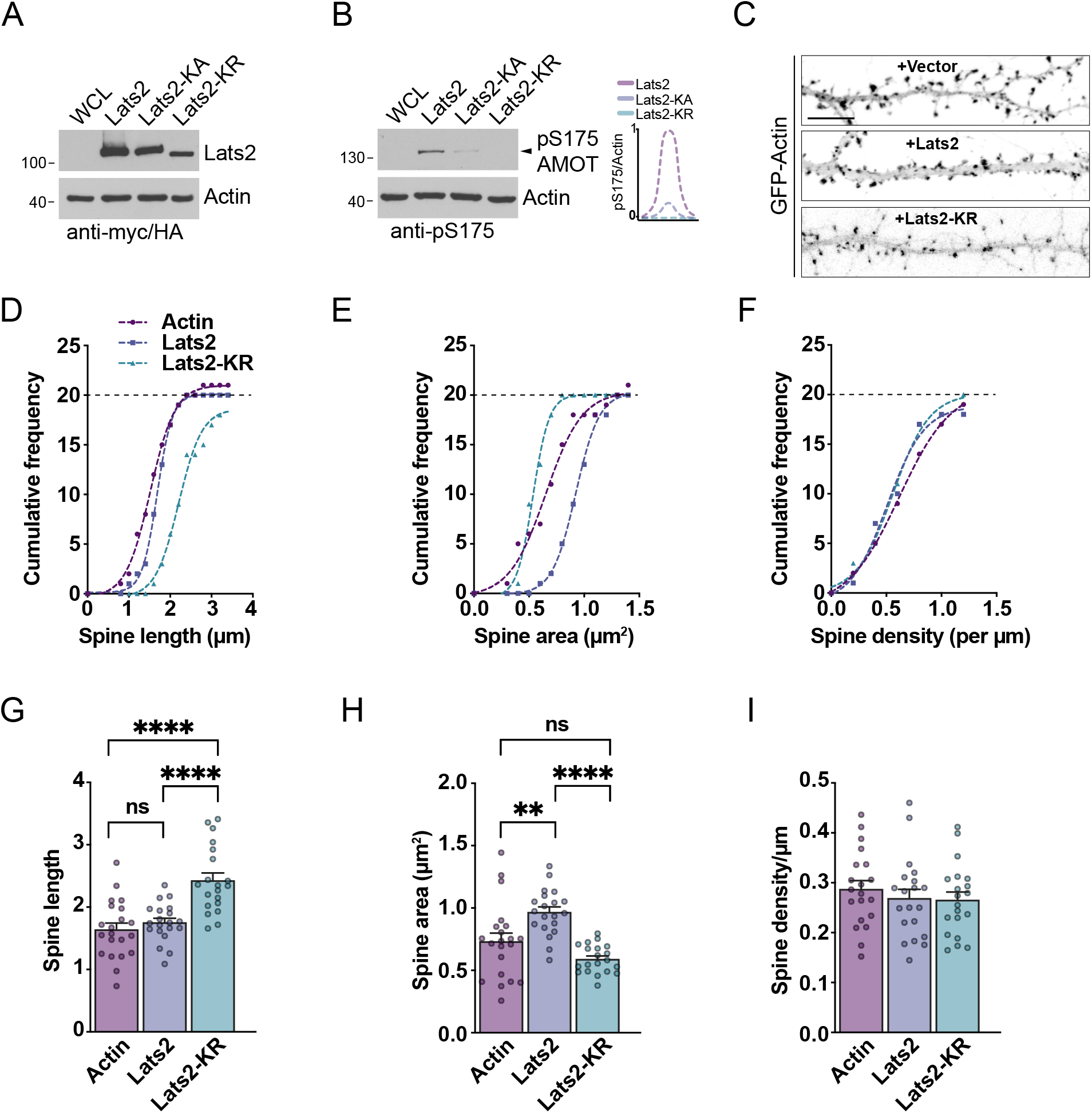
LATS2 affects dendritic spine length and width. **A.** Immunoblot showing expression of myc-LATS2 and kinase-inactive mutants myc-LATS2-KA and HA-LATS2-KR in HEK293 cells. Membranes were probed with anti-myc and anti-HA antibodies and re-probed with anti-actin antibody. **B.** Immunoblot showing phosphorylation of AMOT-130 at serine 175 (pS175) by wild-type (WT) LATS2, but not by the kinase-inactive mutants LATS2-KA and LATS2-KR. Membranes were re-probed with anti-actin antibody. Line scan showing the pS175 signal normalized to actin. **C.** Neurons transfected with GFP-Actin together with empty vector or plasmids expressing myc-LATS2 or HA-LATS2-KR. Scale bar, 10 µm. **D**-**F**. Cumulative frequency distribution of dendritic spine area, spine length, and spine density in neurons co-expressing GFP-Actin and myc-LATS2 or HA-LATS2-KR. **G**-**I**. Quantification of dendritic spine length, spine area and spine density in neurons co-expressing GFP-actin and WT LATS2 or LATS2-KR. Data represent means ± SEM. \*\*\*\**P* < 0.0001; *P* = 0.7009 (ns), one-way Anova with Tukey’s multiple comparisons test.

We next examined presynaptic organization by analyzing Bassoon localization in transfected neurons (**Fig. 4A-C**). Both WT and kinase-dead LATS2 showed a reduction in Bassoon puncta per unit dendrite length compared to actin alone with no difference detected between the two constructs (**Fig. 4D**; \*\**P* < 0.01; \*\*\**P* < 0.001, Anova with Tukey’s multiple comparison). As spine density was unchanged (**Fig. 3I**), this reduction suggests altered synaptic organization rather than structural elimination of spines. Despite similar overall puncta density between WT and kinase-dead LATS2, the subcellular Bassoon localization differed markedly. WT LATS2 exhibited greater number of Bassoon puncta apposed to spine heads relative to dendritic shafts (**Fig. 4E**; \*\**P* < 0.01; \*\*\*\**P* < 0.001, Anova with Tukey’s multiple comparison), whereas LATS2-KR showed the opposite pattern, with most Bassoon puncta distributed along dendrites (**Fig. 4E**; \*\**P* < 0.01; \*\*\*\**P* < 0.001, Anova with Tukey’s multiple comparison). This redistribution likely reflects immature protrusion formation and suggests that LATS2, either directly or via postsynaptic interactions, destabilizes synaptic adhesion without affecting overall protrusion density.

**Figure 4.**
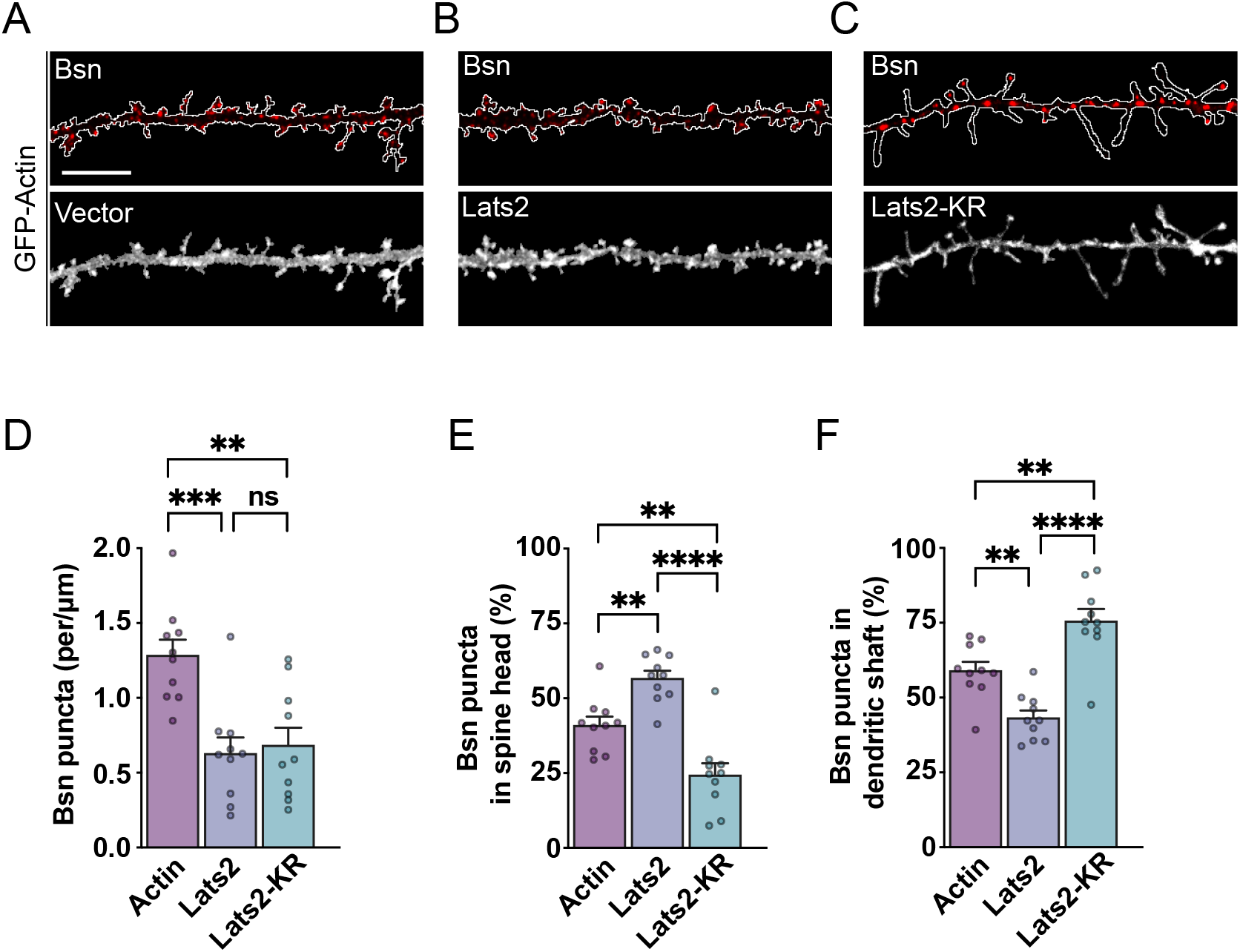
LATS2 kinase activity affects synapse organization. **A**-**C**. DIV 16 neurons expressing GFP-Actin, myc-LATS2 or HA-LATS2-KR were immunostained for endogenous Bassoon (Bsn); LATS2 and LATS2-KR were detected using anti-myc and anti-HA antibody, respectively. Scale bar, 10 µm. **D.** Total numbers of bassoon puncta per µm dendrite length. Data represent means ± SEM. \*\*\**P* < 0.001; \*\**P* < 0.01; *P* = 0.934 (ns), one-way Anova with Tukey’s multiple comparisons test. **E.** Fraction of bassoon puncta in spine heads. Data represent means ± SEM. \*\*\*\**P* < 0.0001; \*\**P* < 0.01, one-way Anova with Tukey’s multiple comparisons test. **F.** Fraction of bassoon puncta in dendritic shafts. Data represent means ± SEM. \*\*\*\**P* < 0.0001; \*\**P* < 0.01, one-way Anova with Tukey’s multiple comparisons test.

LATS2 contains a kinase domain located in the C-terminal region and an N-terminal unstructured region known to mediate binding to signaling proteins (Furth and Aylon, 2017). To determine the region responsible for LATS2 spine localization we generated myc-tagged deletion constructs encoding the N-terminal region (ΔC; 1-720), and the C-terminal kinase-containing region (ΔN; 667-1088) (**Fig. 5A**). Expression of the constructs was confirmed in COS-7 cells and verified by immunoblotting using anti–myc antibody (**Fig. 5B**). We next validated their subcellular distribution in neurons, which revealed a graded enrichment pattern with the N-terminal construct showing the highest enrichment in spine heads, the full-length protein exhibiting intermediate enrichment, and the C-terminal construct localizing predominantly along the dendrite (**Fig. 5C**). To quantify the accumulation of each LATS2 construct at synapses, we calculated the spine enrichment factor (E_f_) by normalizing spine myc-tag fluorescence to tdT in dendritic shafts as previously described (Wigerius et al., 2018). This revealed that the N-terminal construct enriched 50 % more in spine heads than full-length LATS2, whereas the C-terminal construct showed ∼ 30 % less enrichment relative to full-length protein (**Fig. 5D**; \*\*\**P* < 0.001; \*\**P* < 0.01; *P* = 0.1601 (ns), one-way Anova with Tukey’s multiple comparisons test). As LATS2-ΔN which lack the N-terminal region failed to localize in spines, these findings ascertain that the N-terminal region is necessary for the localization of LATS2 in dendritic spines. They also suggest that postsynaptic proteins might bind LATS2 within this region contributing to its compartment-specific localization.

**Figure 5.**
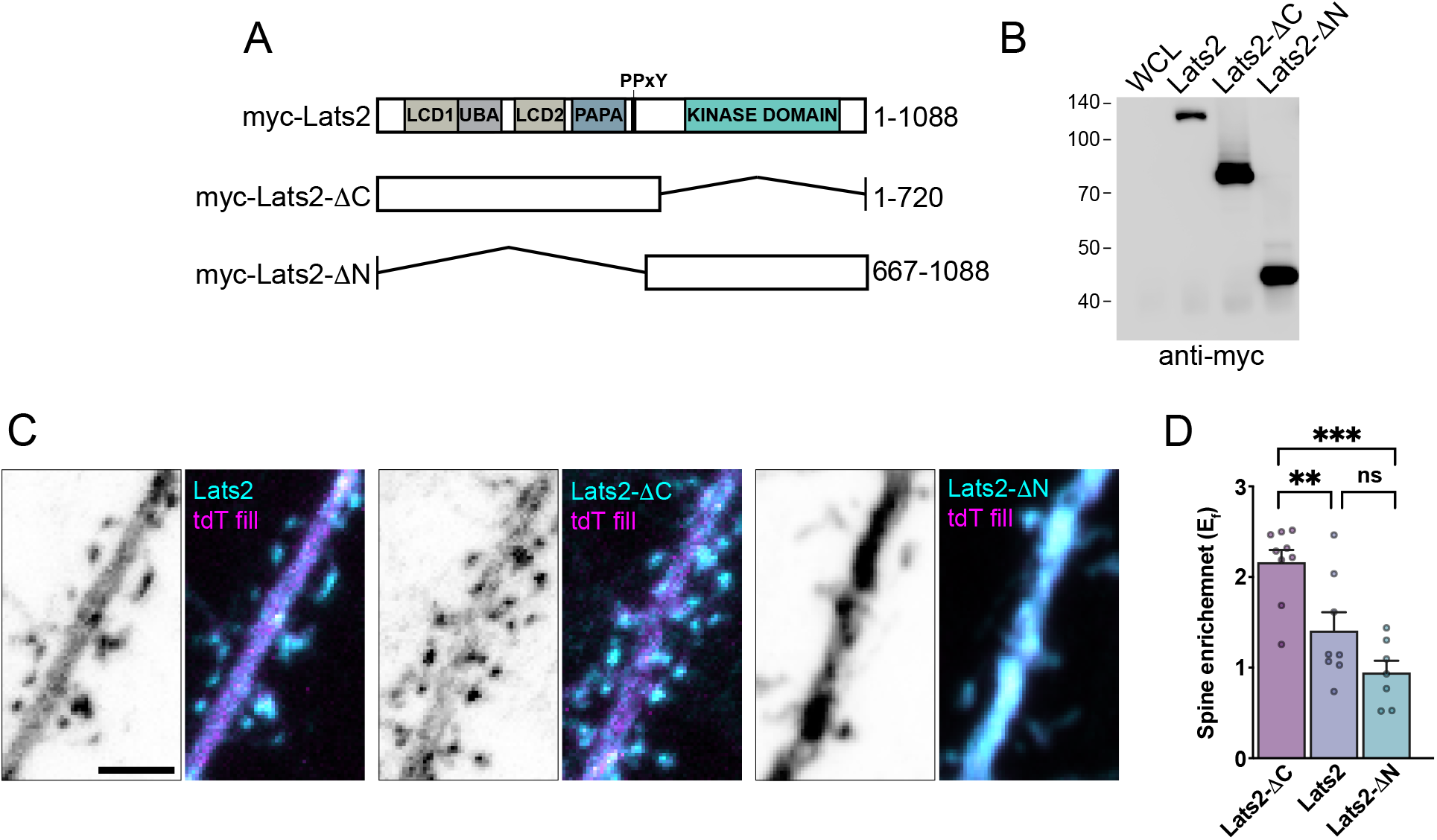
The LATS2 N-terminal mediates spine enrichment. **A.** Schematic showing LATS2 domains and constructs used. **B.** Immunoblot of myc-LATS2, myc-LATS2-ΔC, and myc-LATS2-ΔN expression in HEK293 cells. Membranes were probed with anti-myc antibody. Whole cell lysate (WCL). **C.** DIV 15 neurons co-expressing a fill, tandem dimer tomato (tdT), together with myc-LATS2-FL (1-1088), myc-LATS2-ΔC (1-720), or myc-LATS2-ΔN (667-1088) were immunostained using antibody against the myc-tag. Panels show dendritic segments with expression in spine heads and shafts. Scale bar, 3 µm. **D.** Spine head enrichment (E_f_) of the indicated LATS2 constructs (cyan) relative to tdT (magenta) was calculated as described in methods. Data represent means ± SEM. \*\*\**P* < 0.001; \*\**P* < 0.01; *P* = 0.160 (ns), one-way Anova with Tukey’s multiple comparisons test.

LATS1 has previously been reported to modulate actin filaments (Visser-Grieve et al., 2011), the main structural components in neurons. Given that LATS2 causes changes in the spine structure, we examined whether these changes are linked to actin dynamics. We first tested whether LATS2 forms a complex with Actin by performing immunoprecipitation using a GFP antibody in lysates from HEK293 cells co-expressing GFP-Actin and myc-LATS2, myc-LATS2-ΔC, or myc-LATS2-ΔN. Both full-length LATS2 and LATS2-ΔC associated with actin, with the N-terminal region showing strong enrichment in the complex (**Fig. 6A**), arguing that this region is necessary for binding to actin. In contrast, LATS2-ΔN containing the kinase domain failed to form a detectable complex. We next analyzed the distribution of GFP-Actin and myc-LATS2 in 15-16 DIV hippocampal neurons, which demonstrated significant accumulation of clusters in neurons (**Fig. 6B**). Analyses of fluorescence intensities revealed that both proteins overlapped primarily in spine heads and less in dendritic shafts (**Fig. 6C**; spine heads: *r* = 0.823 and shafts: *r* = 0.311; *Z* = 5.90, *\*\*\*\*P* < 0.0001, two-tailed Fisher’s r-to-z test). To more directly determine how LATS2 affects actin we examined the turnover of GFP-Actin in spines by performing fluorescence recovery after photobleaching (FRAP) experiments. The mobility of GFP-Actin, co-expressed together with myc-LATS2 or HA-LATS2-KR mutant, was tested 2 days after transfection (**Fig. 6D-F**). We found that myc-LATS2 reduced GFP-Actin mobility to 17 ± 6.44 % whereas in HA-LATS2-KR expressing neurons the mobile fraction was 58.05 ± 9.38%, approximately 1.5-fold higher than in WT LATS2 expressing neurons, but still lower than with GFP-Actin alone at 85.96 ± 3.37% (**Fig. 6E**, **6F**; *\*\*\*\*P* < 0.0001; \*\**P* < 0.01, one-way Anova with Tukey’s multiple comparisons test). These results suggest that LATS2 affects spine structure by targeting actin filaments. Moreover, although actin turnover differed between WT and mutant LATS2 is reduced suggesting that the kinase activity maybe coupled to actin cytoskeleton regulation. Future work will help define this.

**Figure 6.**
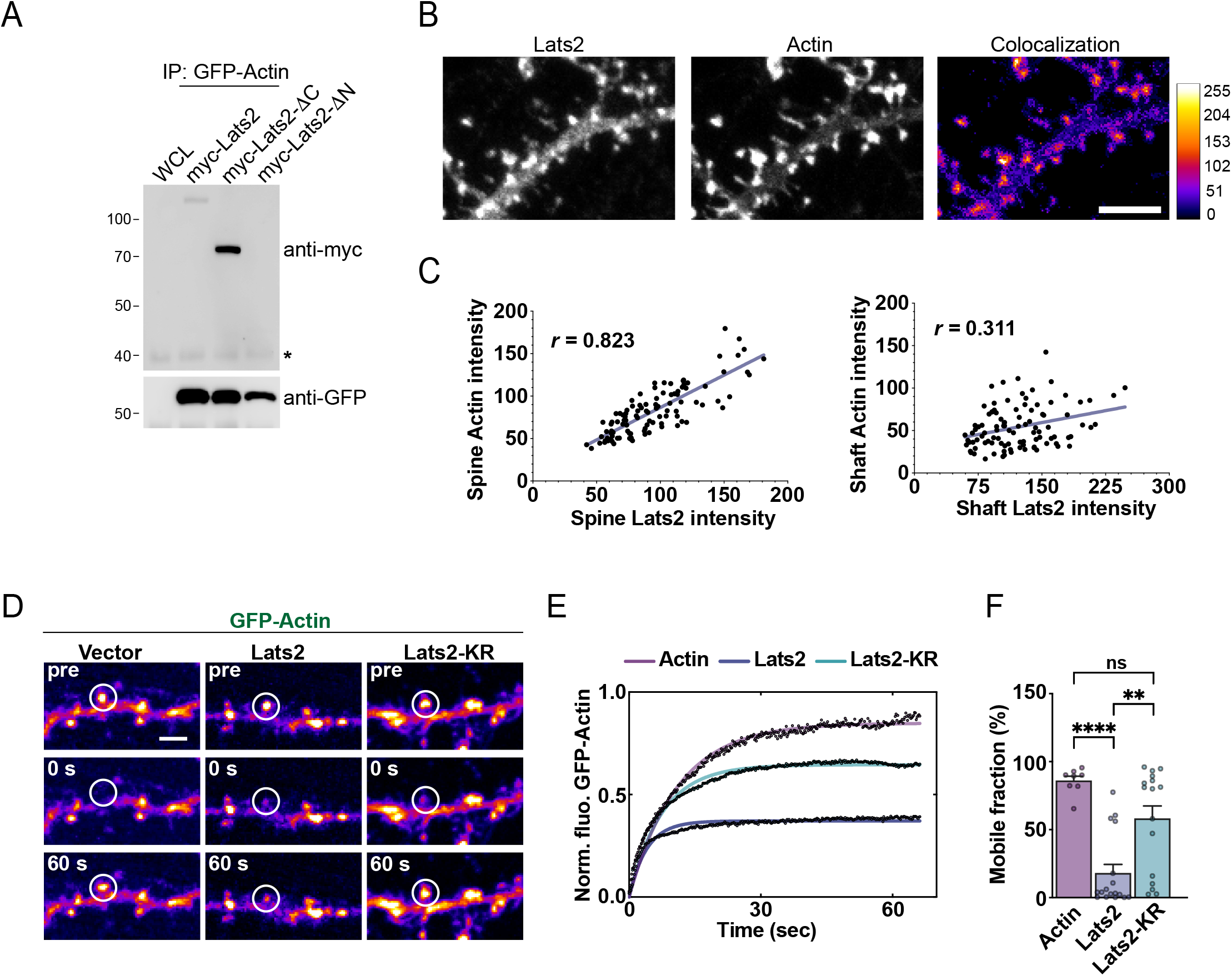
LATS2 regulates GFP-Actin dynamics. **A.** Lysate from HEK293 cells expressing GFP-actin together with myc-LATS2, myc-LATS2-ΔC or myc-LATS2-ΔN were immunoprecipitated with anti-GFP antibody and analyzed by immunoblotting. The membranes were probed with anti-myc recognizing LATS2 or anti-GFP antibody. Negative control (NC). The asterisk indicates the IgG heavy chain. **B.** Neurons co-expressing GFP-Actin and myc-LATS2 were immunostained with anti-myc antibody. Insets show myc-LATS2 (top), GFP-Actin (middle) and their pixel overlap (bottom) with a pseudo colour gradient bar. Scale bar, 5 µm. **C.** Pearson plots of puncta overlap between LATS2 and actin in spine heads (top) and dendritic shafts (bottom): spine heads: *r* = 0.823 and shafts: *r* = 0.311; *Z* = 5.90, *\*\*\*\*P* < 0.0001, two-tailed Fisher’s r-to-z test. **D.** FRAP results showing GFP-Actin alone or expressed together with myc-LATS2 or HA-LATS2-KR. Time-lapse images show spine turnover before and at 30 sec and 60 sec after photobleaching. White circles indicate photobleached spines. **E.** Normalized data fit in single exponential decay curves showing fluorescence recovery (y-axis) plotted against time after bleaching. **F.** Quantification of GFP-Actin recovery in dendritic spines. Data represent mean ± SEM of the mobile fraction after photobleaching. \*\*\*\**P* < 0.0001, \*\**P* < 0.01, *P* = 0.087 (ns), one-way Anova with Tukey’s multiple comparisons test.

LATS2 interacts directly with the scaffolding protein Kibra (Xiao et al., 2011) which forms a complex with AMPARs and facilitates activity-triggered receptor recycling at the plasma membrane (Stepan et al., 2024). Additionally, LATS2 has been reported to regulate the Kibra-AMPAR interaction(Stepan et al., 2024). We therefore tested whether LATS2 is involved in AMPAR recycling at the plasma membrane by co-expressing super-ecliptic pHluorin (SEP) fused to the N-terminus of the glutamate receptor 1 (SEP-GluR1) in neurons. We observed strong overlap between SEP-GluR1 and both endogenous GluR1 (data not shown) and myc-LATS2 (**Fig. 7A**) along dendrites and spines 3 days after transfection, suggesting proper trafficking and postsynaptic distribution of SEP-GluR1. However, there was no discernable difference in puncta overlap between spines and dendritic shafts (**Fig. 7B**; Spines (*r* = 0.744) and shafts: (*r* = 0.833); *Z* = -1.69, *P* = 0.092 (ns), two-tailed Fisher’s r-to-z test). SEP-GluR1 fluorescence intensity increase in response to synaptic activity due to enhanced trafficking of AMPARs at the plasma membrane. To monitor activity-dependent insertion of SEP-GluR1 fluorescence, neuronal cultures were stimulated with 300 µM glycine for 4 minutes. Glycine elicited a linear increase in fluorescence over time in spines expressing SEP-GluR1 alone (**Fig. 7C**). This increase was reduced by approximately 2.5-fold in spines overexpressing myc-LATS2 (**Fig. 7D**) and was evident at both 10 minutes (SEP-GluR1: 1.14 vs SEP-GluR1 + LATS2: 0.45, \**P* < 0.05) and 20 minutes (1.60 vs 0.66, \*\**P* < 0.01), with no significant difference between the two groups prior to treatment (0.74 vs o.61, *P* = 0.93; two-way Anova with Sidak’s multiple comparisons test). Taken together, these findings support a role for LATS2 in AMPAR trafficking consistent with its link to Kibra in dendritic spines.

**Figure 7.**
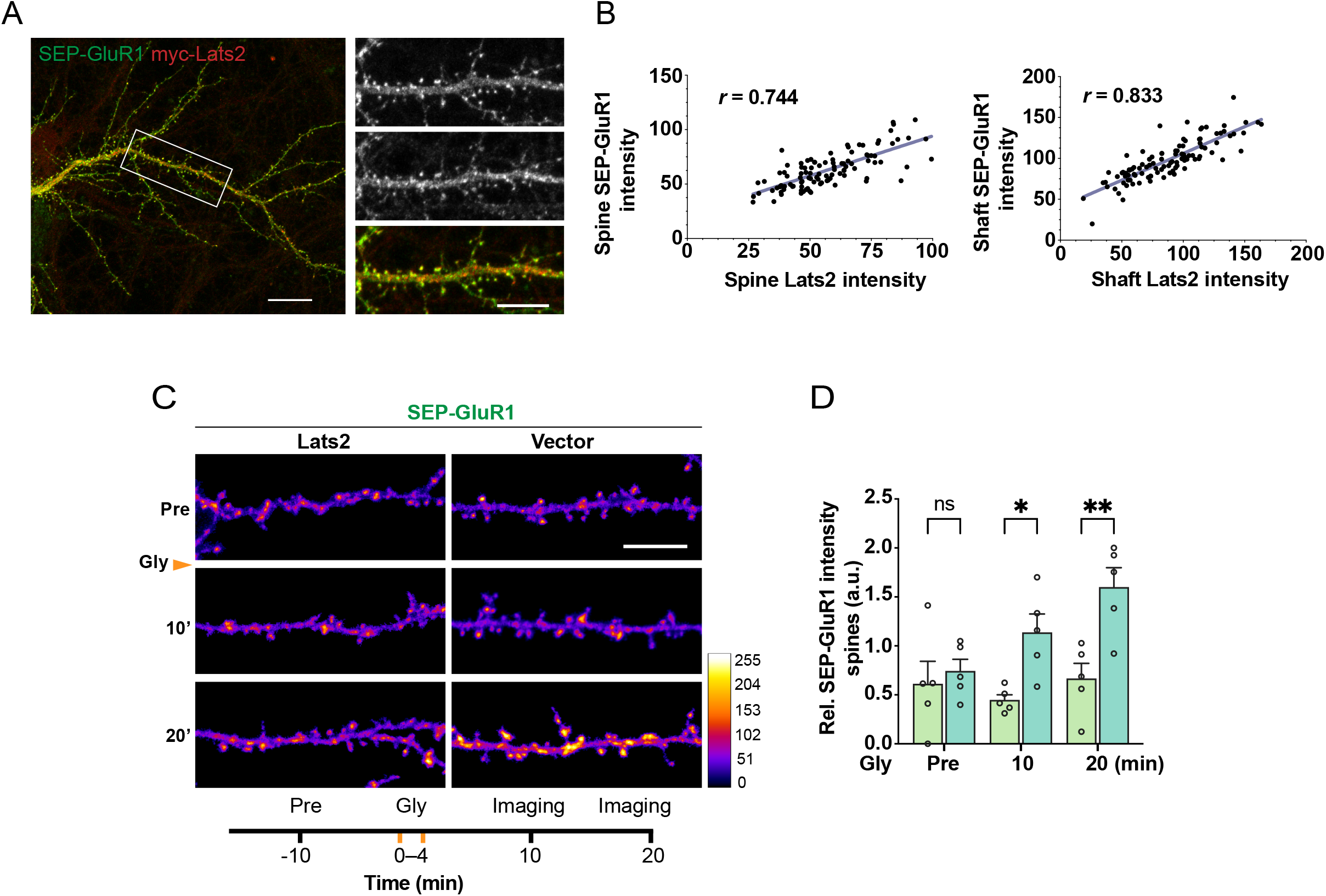
LATS2 suppresses SEP-GluR1 fluorescence. **A.** Neurons co-expressing myc-LATS2 and SEP-GluR1 were immunostained with anti-myc antibody. Magnified insets to the right show expression of SEP-GluR1, myc-LATS2 and the overlay. Scale bars, 20 µm and 10 µm. **B.** Pearson plots of puncta overlap between myc-LATS2 and SEP-GluR1 in spine heads and dendritic shafts: spine heads: (*r* = 0.744) and shafts: (*r* = 0.833); *Z* = -1.69, *P* = 0.092 (ns), two-tailed Fisher’s r-to-z test. **C.** Representative images of 16 DIV dendrites expressing SEP-GluR1 together with vector or myc-LATS2 stained with anti-myc antibody. Neurons were imaged before and at 10 and 30 minutes after treatment with 200 µM glycine (Gly) for 4 minutes. SEP-GluR1 intensity is pseudo coloured with gradient shown. Scale bar, 10 µm. **D.** Quantification of SEP-GluR1 fluorescence intensity at the indicated time points. Data represent mean ± SEM. \*\**P* < 0.01, \**P* < 0.05, *P* = 0.932 (ns), two-way Anova with Sidak’s multiple comparisons test.

## Discussion

Our data suggest that LATS2 influences dendritic spine maturity by controlling actin dynamics. While LATS2, which associates with F-actin, impacts spine size, expression of a dominant-negative (DN) LATS2 mutant with reduced catalytic activity trigger formation of immature protrusions and a disruption of synaptic bassoon clusters. These effects correlated with reduced SEP-GluR1 fluorescence in neurons co-expressing LATS2 after brief glycine stimulation. Thus, LATS2 expression causes spine instability and aberrant shape which leads to inability to form proper synaptic connections. These effects are likely caused by loss of actin dynamics which disrupts spine structure, driving synaptic AMPARs removal at the postsynaptic membrane. Thus, consistent with our earlier findings on LATS1 and the current study on LATS2, we conclude that both proteins function through synaptic F-actin although the mechanisms by which they operate differs.

LATS1 and LATS2 are encoded by different genes and regulated by different gene-regulatory networks (Furth and Aylon, 2017). Consistent with this, we found that *LATS2* mRNA levels in mice increase across development, with higher expression in the adult brain compared to the prenatal brain, which is tightly regulated during development (**Fig. 1**). We also observed elevated LATS2 protein in neuronal cultures at DIV4 with levels remaining steady at ensuing time points (**Fig. 2**). This is in contrast to LATS1 whose expression diminishes around DIV7 and remains undetectable at later stages (Wigerius et al., 2018). Human brain transcriptomics data further show that *LATS2* is highly enriched in the CA3/CA4 regions, which are functionally involved in processing memory-based behaviours in the hippocampus (Hainmueller and Bartos, 2020). Neurons in these regions are characterized by large synapses that make contact simultaneously with multiple dendritic spines. LATS kinases may act synergistically to establish these contacts, and once stable circuits have formed in mature neurons, it is likely LATS2 compensates for the absence of LATS1.

Expression of a catalytic-inactive LATS2 mutant increases spine length while WT LATS2 enhances spine head size (**Fig. 3**). These effects are likely the result of changes in actin cytoskeleton remodeling, consistent with the idea that LATS2 affects either the concentration or function of trans-synaptic regulators, although we don’t know yet whether this occurs through direct phosphorylation or via a scaffolding role. Moreover, we found a decrease in the levels of apposing presynaptic bassoon puncta in neurons expressing the LATS2 mutant (**Fig. 4**). Since spine density was unaffected in these neurons, we interpret this to be the result of the overall disturbed adhesion between synaptic contacts. In neurons, Ndr1/2 kinases have been reported to exert similar effects on synaptic integrity (Ultanir et al., 2012; Sharif and Hergovich, 2018), indicating that Hippo kinases participate in broadly conserved signaling pathways within the brain.

In this study we found that the N-terminal in LATS2, which in other systems mediates binding to various components including the scaffolding protein Kibra (Xiao et al., 2011), is required for spine head enrichment (**Fig. 5**). This result is consistent with a widely accepted role for the region as a targeting platform that enable scaffolding. Indeed, we detected an association with actin mediated via the N-terminal and both proteins showed significant puncta overlap in spine heads (**Fig. 6**) providing further evidence that the region is central to the subcellular synaptic localization of LATS2. The N-terminal is also specifically responsible for the interaction between LATS1 and actin in fibroblasts (Visser-Grieve et al., 2011), although the mechanism by which the two LATS kinases interact and influence cell behaviour may differ. We found that LATS2 over-expression increases spine head size, an effect typically associated with enhanced actin polymerization, while LATS1 was reported to inhibit this process in mouse fibroblasts independently of its kinase activity. However, the catalytic-inactive mutant produces immature, long protrusions suggesting that the LATS2 kinase domain carry information that contributes to its effect on actin dynamics. Thus, it remains unclear exactly how LATS kinases modulate the actin cytoskeleton. One possibility is that they phosphorylate substrates with distinct functions. Alternatively, their relative concentrations may alter cytoskeletal dynamics or levels of actin-binding factors that regulate this process. In FRAP analysis, we observed a significant stabilization of actin filaments by LATS2 that was reversed to some extent by the catalytic-inactive mutant (**Fig. 6**). Since actin filaments are the main cytoskeletal structures responsible for spine shape modulation (Okamoto et al., 2004), these data suggest that abnormal spine morphology by LATS2 is a consequence of disturbed actin dynamics.

What other factors associate with LATS2 in the synapse? Although both LATS kinases have been reported to interact with Kibra in spines and their inactivation results in the release of Kibra (Stepan et al., 2024), the mechanism is not clear. Kibra drives AMPAR subunit membrane assembly and trafficking (Heitz et al., 2016; Tracy et al., 2016), and evidence suggests this occurs only in adult animals lacking Kibra at a stage when its levels are low, raising the possibility that Kibra expression is regulated during development (Makuch et al., 2011). Additionally, no direct interaction between Kibra and AMPARs has been reported. Instead, Kibra influences receptor function via the actin-binding protein PICK1, which binds AMPAR subunits and functions as a master organizer for their assembly and trafficking (Makuch et al., 2011; Hanley, 2018). While we were unable to detect a physical interaction between LATS2 and AMPARs in the rodent brain (data not shown), we found that expression of LATS2 in neurons suppressed SEP-GluR1 insertion into the plasma membrane in response to glycine application (**Fig. 6**). This effect persisted over the monitored time points while control neurons expressing SEP-GluR1 alone underwent a rapid fluorescence intensity increase. While these results do not address the precise mechanism for this observation, they suggest reduced SEP-GluR1 fluorescence might arise from disturbed actin dynamics causing a delay in AMPAR capturing at the postsynaptic membrane. Apart from scaffolding Kibra in the synapse, LATS2 might therefore hold a more direct role in the delivery of AMPARs to the membrane through its association with the actin cytoskeleton. This idea is consistent with the view that actin dynamics is essential for both clustering and insertion of AMPARs at the postsynaptic membrane, while stabilization of actin filaments disrupt this process (Gu et al., 2010; Hanley, 2018). Yet, the elucidation of LATS2-mediated inhibition of AMPAR trafficking require future experiments, and whether LATS2 binds other PSD components, including PICK1, to mediate this effect remains to be determined.

The LATS1/2 kinases participate in multiple signaling pathways and engage in a complex set of interactions that vary across cellular contexts. In some cases, their functions appear redundant through involvement of shared pathways, whereas in others they act independently. Further investigation of LATS2 in the nervous system may uncover novel mechanism and binding partners, while also revealing additional roles for LATS kinases in their canonical regulation of cell polarity and organ growth.

## Acknowledgements

This work was supported by funding from Discovery grants RGPIN-2022-03370 (J.P.F) and RGPIN-2021-03303 (S.R.K) from the National Science and Engineering Council (NSERC) of Canada.

## Conflict of interest statement

The authors declare no competing financial interests.

## Notes

### Competing Interest Statement

The authors have declared no competing interest.

